# Endothelial cell MHC molecules are necessary and sufficient to reject 3D-printed human skin grafts in an advanced human immune system mouse

**DOI:** 10.1101/2025.11.24.689811

**Authors:** Zuzana Tobiasova, Esen Sefik, Lingfeng Qin, Jennifer M. McNiff, Gwendolyn Davis, Richard A. Flavell, W. Mark Saltzman, Jordan S. Pober

**Affiliations:** Department of Immunobiology, Yale University, USA; Department of Surgery, Yale University, USA; Department of Dermatology, Yale University, USA; Department of Pathology, Yale University, USA; HHMI (Howard Hughes Medical Institute, Yale School of Medicine, Yale University, USA; Department of Biomedical Engineering, Yale University, USA; Department of Chemical & Environmental Engineering, Yale University, USA; Department of Cellular & Molecular Physiology, Yale University, USA

## Abstract

Vascularized skins were 3D-printed using single donor human fibroblasts, pericytes, keratinocytes and endothelial cells (ECs), the latter either unmodified (WT-ECs) or deleted of MHC molecules (KO-ECs). Adult MISTRG6 immunodeficient mice neonatally inoculated with adult human hematopoietic stem cells (HSCs) received printed skin allogeneic to the HSCs and were boosted 3 weeks post-grafting with human peripheral blood mononuclear cells (PBMCs) autologous to the HSCs. HSC inoculation alone produced low levels of circulating human myeloid and lymphoid cells without affecting grafts; PBMC boosting dramatically increased circulating human CD4+ T cells and boosted CD8+ T cells only in mice with WT-EC grafts. These grafts became infiltrated by human macrophages, dendritic cells, CD4+ and CD8+ T cells, and showed evidence of rejection. Shared T cell clones were present in skin and spleen. KO-EC grafts had minimal infiltration of graft or spleen without rejection despite MHC molecule expression on other graft cell types.

## INTRODUCTION

Tissue engineering can provide a potentially unlimited source of organs for therapeutic transplantation, but currently are restricted to thin constructs because they lack vascularization and must survive by nutrient and gas diffusion (*1*). Skin grafting is used to treat non-healing ulcers and extensive skin tumor resections. Autologous skin transplantation is limited because the underlying disorders that lead to non-healing ulcers, such as diabetes or advanced age, generally impair wound repair, complicating skin harvest. (*2*) Allogeneic skin is among the tissues most prone to rejection (*3*). This need has driven development of synthetic skin substitutes. Burke and Yannas described clinical results using bilayered artificial skin as early as 1981 (*4*), and there are over ten different approved synthetic skin substitutes that are either acellular or cellularized to form dermal, epidermal, or epidermo-dermal mimetics. (*5, 6*) None are completely satisfactory (*7*).

Cellularized grafts better respond to environmental stresses and injuries, although acellular grafts may become cellularized after implantation through invasion of the graft material by recipient cells. To be clinically useful, grafts must be available “off the shelf.” Consequently, the currently available cellularized skin substitutes contain keratinocyes and/or fibroblasts that are allogeneic to the host. Surprisingly, unlike allogeneic natural skin, allogeneic cells in these synthetic skins do not initiate a recipient immune-mediated rejection response (*8, 9*). Nevertheless, the graft cells die and grafts often slough due to the absence of microvascular graft perfusion (*10*). Incorporating a vascular network into the graft could avoid ischemic sloughing. A critical component of the microvascular circulation is the endothelial cell (EC) lining, which regulates blood flow, controls permselectivity, and minimizes thrombosis and inflammation. (*11*) Moreover, cultured human ECs can spontaneously organize into a tubular network capable of perfusing acellular collagen gels or decellularized human skin (*12, 13*), suggesting that inclusion of ECs in a synthetic graft could provide adequate perfusion in skin and perhaps other tissues (*14*).

The absence of immune rejection of constructs containing allogeneic cells is puzzling, but can be explained in two ways: either grafts lack specific cell types capable of alloantigen presentation to resting T cells, such as myeloid or lymphoid cells resident within normal tissues, or, alternatively, that absence of a vascular system within the graft limits contact between recipient alloreactive T cells and graft cells. Thus including ECs to form blood vessels may address lack of perfusion, it may simultaneously lead to rejection regardless of which explanation is true because human ECs express and can present both class I and II MHC molecules, activating both alloreactive CD4+ and CD8+ effector memory T cells (*15*). Alloreactive T effector memory T cells arise as a result of prior infections because some T cell receptors (TCRs) for antigen specific pathogen peptide bound to self MHC cross react to non-self MHC molecules independent of the bound peptide (*16*). In clinical kidney transplantation, rejection is best predicted by the frequency of pre-existing alloreactive effector memory T cells rather than total alloreactive T cells (*17*), and in immunodeficient mice bearing human skin grafts, adoptively transferred human effector memory T cells are the only T cell subpopulation capable of causing acute cell-mediated rejection (*18*). Much of the acute rejection response is mediated by CD8+ effector T cells, also known as cytotoxic T lymphocytes (CTLs), that directly recognize graft class I MHC molecules, but activated host CD4+ T cells can boost the CD8+ T cell response or activate natural killer cells or macrophages (*19*). Activation of host CD8+ or CD4+ T cells may involve recognition of non-self class I or II MHC molecules, respectively, on a graft-derived ECs (called “direct recognition”), or on a host myeloid or lymphoid antigen presenting cell that has acquired intact MHC molecules shed from the graft (called “semidirect recognition”). A third pathway (“indirect recognition”) involves host antigen presenting cells acquiring peptides from allogeneic graft proteins and presenting them to host CD4+ T cells, but this process typically occurs with some delay. CRISPR-mediated gene disruption can be used to eliminate the expression of both class I and II MHC molecules on ECs, ablating their capacity to directly initiate an allogeneic response without compromising their capacity to self-assemble into blood vessels (*20*). Microvessels formed from such “knock out ECs” (KO-ECs) could still promote contact of T cells with graft cells but could no longer act as antigen presenting cells (APCs).

We and others have studied the process of skin graft rejection using immunodeficient mouse strains in which the human immune system is partly reconstituted using hematopoietic cells from a donor allogeneic to the skin (*12*). The majority of human immune system mouse models, including the ones we have used in the past, are deficient in reconstitution of human myeloid cells (*21*) and may be of limited use in assessing rejection initiated by indirect or semi-direct recognition. The MISTRG6 mouse—constructed by incorporating several human genes that support hematopoietic development into an immunodeficient Balb/c-derived strain lacking both *Rag2* and the cytokine common gamma chain (*Il2rg*^-/-^)—was developed to provide conditions that allow both human lymphoid and myeloid cells to be engrafted (*22, 23*). When these animals are injected as neonates with human hematopoietic stem cells (HSCs) isolated from human fetuses, neonates or adults, they develop robust and functional circulating populations of myeloid monocyte/macrophages and dendritic cells in addition to T and B cells. However, these animals are generally raised in specific pathogen-free environments and thus lack the history of infections required to generate alloreactive T memory populations, the principal orchestrators of acute cell-mediated allograft rejection. For this study, we developed a novel use of the MISTRG6 mouse strain in which neonates are inoculated with mobilized adult human HSCs and then engrafted with 3D-printed vascularized human skin grafts formed with a single donor source of keratinocytes, dermal fibroblasts, pericytes and either MHC-expressing (WT) or MHC-deleted (KO) ECs. Three weeks later, the animals are boosted by injection of PBMCs from the same adult donor as the HSCs, thereby engrafting alloreactive effector memory T cells in an animal with functional human myeloid antigen presenting cells (**Figure 1**). We used this model to study the human allogenic immune response to 3D-printed human skin grafts made with either unmodified (WT) ECs or ECs in which both class I and class II MHC molecules have been deleted, allowing us to separate the effects of increased vascularization from the immunological role of ECs.

**Fig 1.**
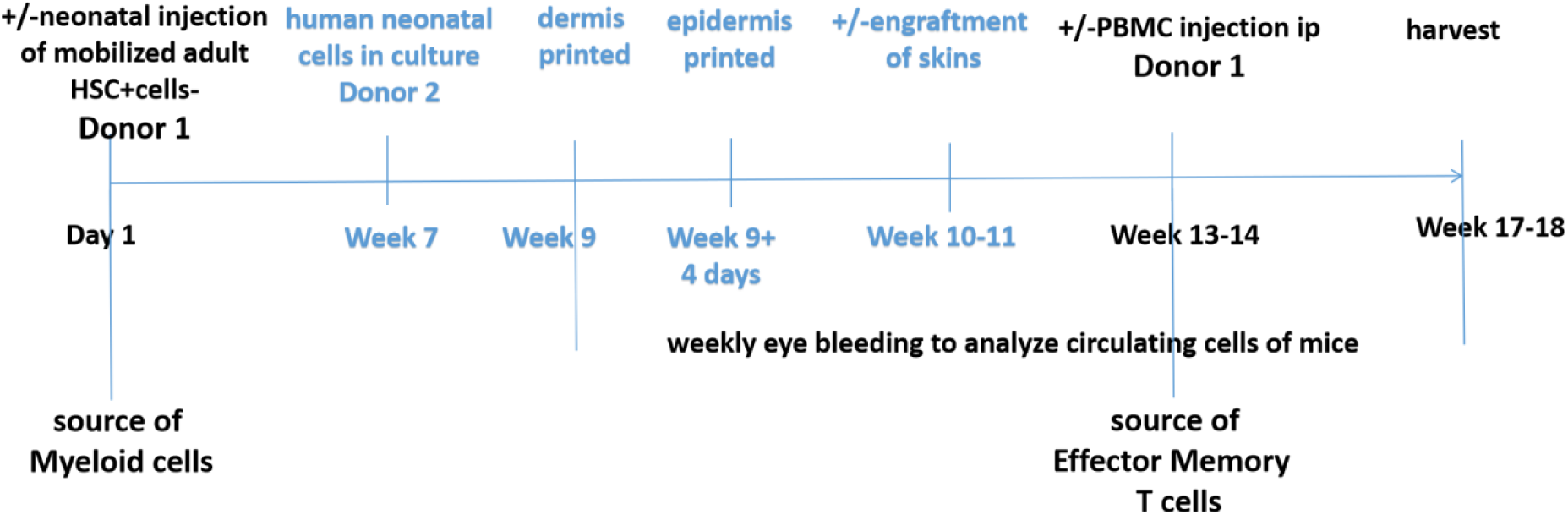
Experimental approach. Timelines of human HSC inoculation, 3D-printed skin grafting and PBMC boosting using MISTRG6 mice.

## RESULTS

### Hematopoietic reconstitution of skin-engrafted mice and 3D-bioprinted skin grafts in vivo

The goal of our study was to determine the potential consequences of vascularizing bioengineered human skin using self-assembly of human ECs to form microvessels, with pericytes and fibroblasts provided as supportive elements. As described in the methods, all cells in the skin construct were from the same donor and the ECs were either unmodified (WT-EC) or altered using CRISPR/Cas9 to ablate expression of both class I and class II HLA molecules (KO-EC). This allowed us to separate the roles of microvascular ECs as a source of perfusion and as a potential initiator of graft rejection. To create our model, we introduced several new elements into the use of human immune system mice to study transplantation.

Although MISTRG6 mice inoculated as neonates with human HSCs develop functional human myeloid as well as lymphoid cells (*24*), these animals do not develop human alloreactive memory T cells when raised specific pathogen free animal care facilities. To address this, we combined neonatal inoculation with human HSCs and boosting as adult animals with PBMCs, a source of alloreactive memory, from the same donor. Each new component was tested in preliminary experiments described in the Methods. Our final experimental scheme is summarized in **Figure 1**. Mice were sacrificed approximately 6-7 weeks after skin engraftment at which time blood, grafts and spleen were examined by histology and flow cytometry. We did not see significant variation among litters or between sex of the mice or between results with different skin cell donors (all male of necessity since foreskin supplies two of the cell types utilized).

We began with analysis of the blood. Mice inoculated as neonates with HSCs and boosted with autologous PBMCs as adults showed a significant expansion of circulating CD45+ human cells that did not occur in mice treated with either HSC or PBMC inoculation alone. The increase in circulating human cells in animals inoculated with HSCs and boosted with PBMCs is comparable to the increased levels in mice that had been implanted with WT-EC or KO-EC skin before receiving the PBMC boost. In several cases if reconstitution was delayed and below 10% at week 3 after PBMC injection, week 4/ harvest data were added for both HSC+WT-EC skin+PBMC and HSC+KO-EC skin+PBMC groups and graph presented as pooled data (**Figure 2A, B**; CD45:HSC+skin, n=12, 3.21±2.02 vs HSC+PBMC, n=3, 35.87±7.36, p<0.0001, HSC+skin vs HSC+WT-EC skin+PBMC, n=11, 34.79±25.41, p=0.0003, HSC+skin vs HSC+PBMC+KO-EC skin, n=4, 24.73±11.73, p<0.0001, CD3: HSC+skin, n=9, 13.55±25.07 vs HSC+PBMC, n=3, 12258.66±4389.84, p<0.0001, HSC+skin vs HSC+PBMC+WT-EC skin, n=11, 5234.13±4310.6, p=0.002, HSC+skin vs HSC+PBMC+KO-EC skin, n=4, 4348.75±4751.22, p=0.01, HSC+PBMC vs HSC+PBMC+WT-EC skin p=0.02). Most of these cells in HSC+PBMC or HSC+WT-EC skin animals were CD4+ T cells (**Figure 2 C, D**). The most striking finding is that the T cell expansion in the skin of mice bearing grafts with WT-ECs but not in mice bearing grafts made with KO-ECs additionally involved a large increase in effector or effector memory (CCR7 negative) CD8+ T cells expressing Granzyme B, typically characterized as CTL or pre-CTL (**Figure 2 C, D; CD4**: HSC+skin, n=7, 1540.43±1433.12 vs HSC+PBMC, n=3, 7079.66±1879.57, p=0.0009, HSC+skin vs HSC+PBMC+WT-EC skin, n=8, 9382±6024.24, p=0.019, HSC+skin vs HSC+PBMC+KO-EC skin n=3, 12367±3340.37, p=0.0005; **CD8**: HSC+skin, n=3, 222.14±387.309, vs HSC+PBMC, n=3, 1468.67±1068.6, p=0.02, HSC+skin vs HSC+PBMC+WT-EC skin, n=8, 13483.87±10894.9, p=0.0069 and HSC+skin vs HSC+PBMC+KO-EC skin, n=3,1234±492.05, p=0.0078, **CD3+CCR7-**: HSC+skin, n=7, 387.43±471.84 vs HSC+PBMC, n=3, 1747.33±541, p=0.0038, HSC+skin vs HSC+PBMC+WT-EC skin, n=5, 14509.6±7933.36, p=0.0007 or HSC+PBMC vs HSC+PBMC+WT-EC skin p=0.035, HSC+PBMC vs HSC+ PBMC+KO-EC skin, n=3, 328.67±445.5, p=0.024, HSC+PBMC+WT-EC skin vs HSC+PBMC+KO-ECskin p=0.002).

**Fig. 2.**
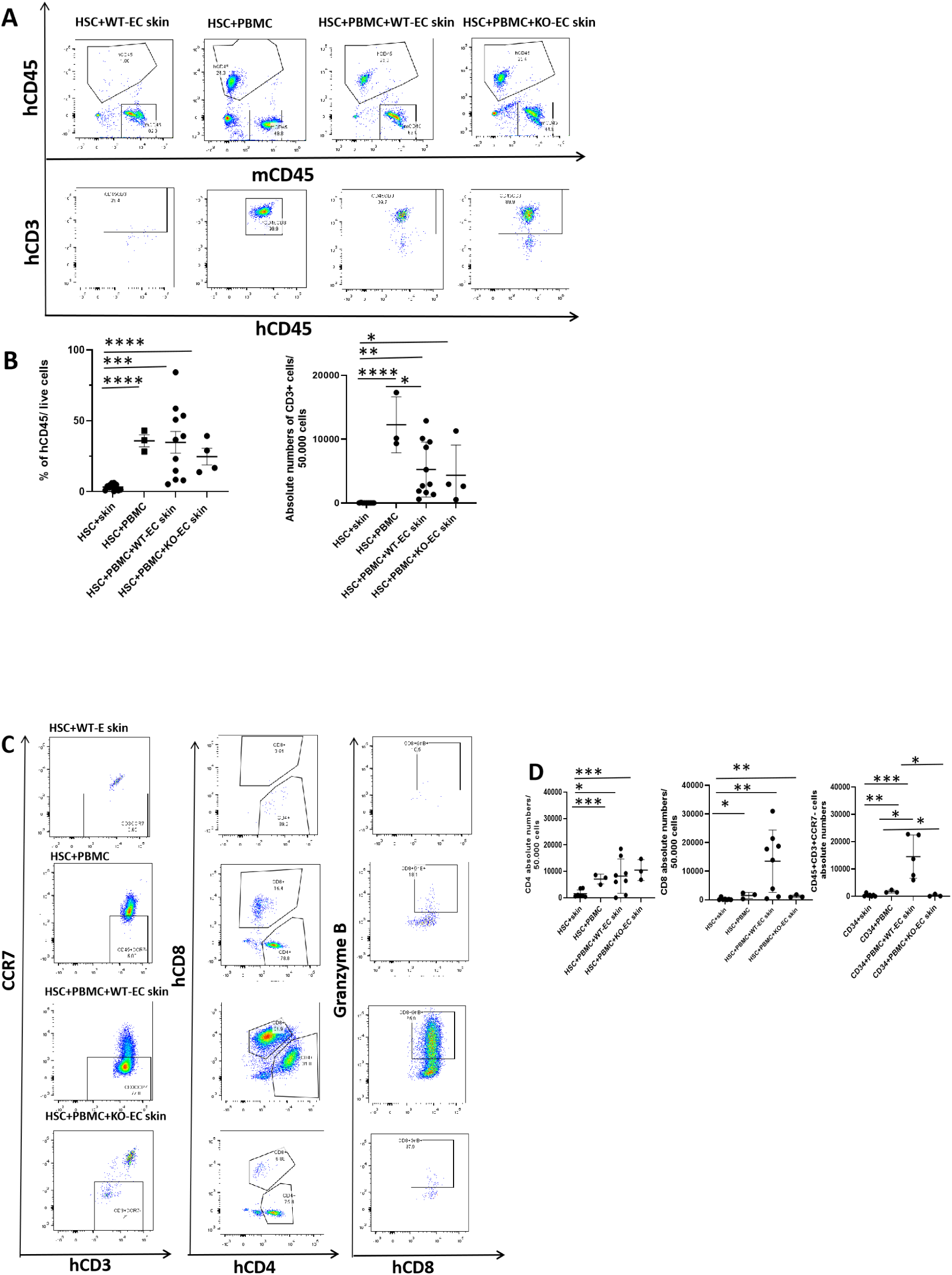
Circulating human cells in skin grafted MISTRG6 mice. **A)** Representative analyses of circulating human leukocytes identified by hCD45 and hCD3 staining in animals inoculated as neonates with HSCs, grafted with synthetic skin and then boosted or not with PBMC assessed 3-4 weeks after boosting. **B)** Quantification of circulating human leukocytes at week 3 after PBMC boost showing increased levels of CD45+ cells in both skin-implanted groups bearing WT-EC skin or KO-EC skin. Absolute numbers of circulating CD3+ lymphocytes show increase after PBMC injection in HSC+PBMC, WT-EC skin and KO-EC skin animals. **C)** Representative phenotyping of circulating CD3+ cells. Majority of expanded T cells in WT-EC skin mice grafts lost CCR7 (effector or effector memory cells), and many express CD8 and granzyme B, indicative of CTL. Remarkably, fewer CCR7- or CD8+ cells are present in KO-EC skin grafts mice. **D)** Quantitation of lymphocytes in circulation at 3-4 week time point after PBMC boost. CD4 numbers show significant elevation in both HSC+WT-EC skin+PBMC and HSC+KO-EC skin+PBMC groups compared to HSC+ skin mice without PBMC boosting. CD8 numbers show significant expansion in HSC+PBMC+WT-EC skin animals in comparison to HSC+skin, HSC+PBMC no skin or HSC+PBMC+KO-EC skin animals. CD3+CCR7-numbers also show significant increase in HSC+PBMC+WT-EC skin animals as compared to HSC+skin or HSC+ PBMC+KO-EC skin mice.

We next examined the bioprinted skin grafts. All of the skin grafts showed HLA-B positive cells, both indicating their human origin and confirming that grafts constructed with KO-EC still expressed HLA antigens on other cell populations (**Figure 3A**). We also found a comparable level of human CD31-lined vessels in both types of grafts (**Figure 3 B, C;** HSC+WT-EC-skin (n=4, 36±9.41), WT-EC-skin+PBMC (n=4, 41.75±16.74) or KO-EC-skin+PBMC (n=3, 49.33±5.13), confirming that KO-ECs retained their capacity to vascularize the constructs. Both WT-EC and KO-EC skin grafts were also infiltrated by mouse EC-lined vessels (**Supplementary Figure 1 A**). Also, skin grafts made with WT-ECs were infiltrated by human leukocytes within 3 to 4 weeks after the boost, largely in a perivascular distribution, and showed histologic evidence of injury. In contrast, skin grafts made with KO-ECs showed minimal evidence of injury (**Figure 3 D-E**; HSC+skin, n=9, 0.05±0.16 vs HSC+PBMC+WT-EC skin, n=12, 1.79±0.7, p<0.0001 and HSC+PBMC+WT-EC skin vs HSC+PBMC+KO-EC skin, n=5, 0.2±0.27 p=0.0006) and developed only minimal levels of infiltration by human CD45RO lymphocytes (**Figure 4 A, B;** HSC+skin, n=4, 0±0 vs HSC+PBMC+WT-EC skin, n=8, 221.25±131.49, p=0.0082, HSC+WT-EC skin vs HSC+PBMC+KO-EC skin, n=3, 33.67±33.5, p=0.04). Human ECs in the WT-EC rejecting grafts showed upregulation of E-selectin, consistent with a human cytokine response but lacked expression in KO-EC skins (**Supplementary Figure 1 B**).

**Fig. 3.**
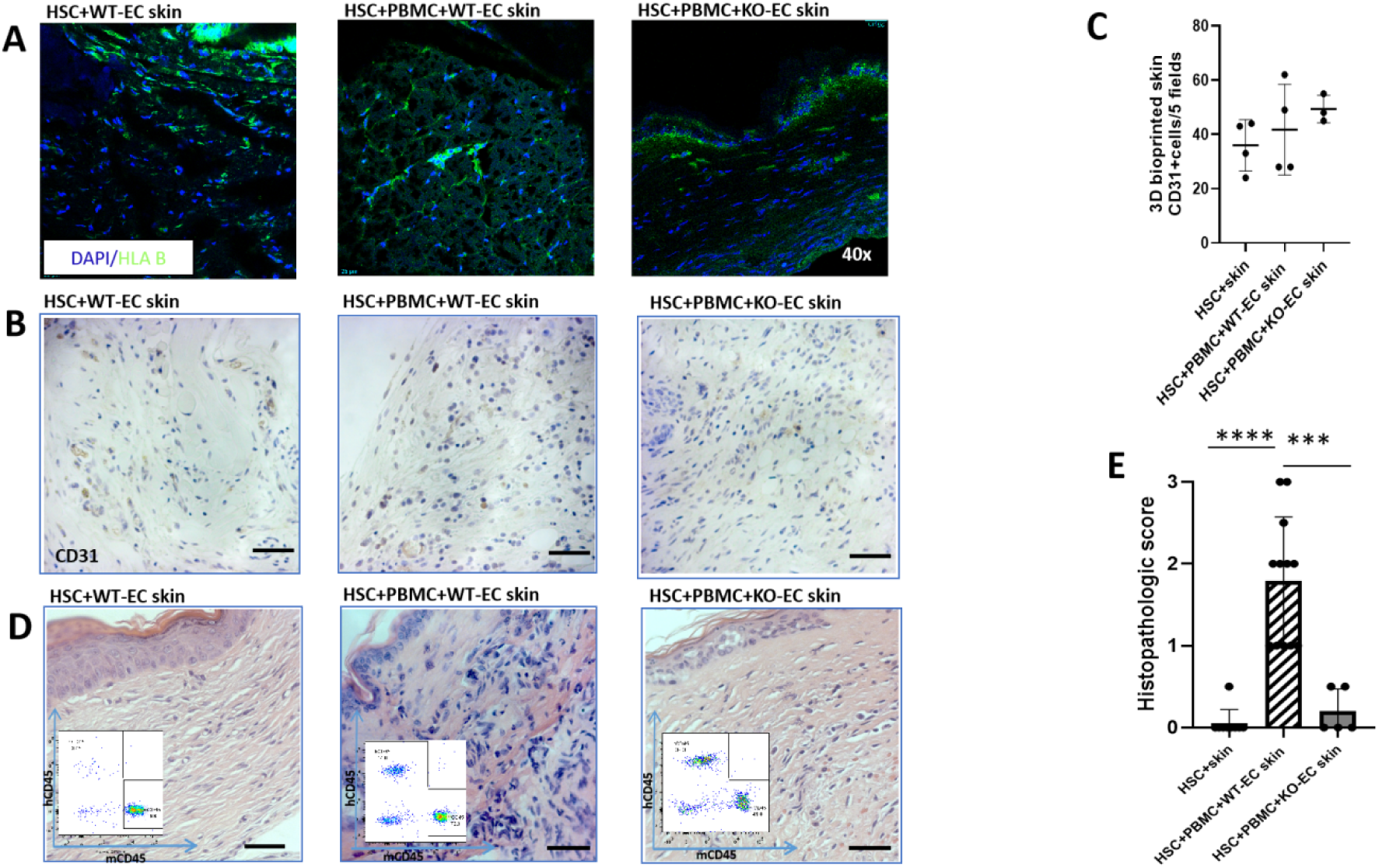
Vitality, vascularization and rejection assessment of 3D-printed skin grafts. **A)** Skin grafts cells presenting HLA B molecules signalizing vital cells present in both HSC+WT-EC skin and HSC+PBMC+KO-EC skin grafts (although not on KO-ECs, still present on other human cells of skin graft). **B,C**) IHC staining of CD31 shows comparable numbers of human ECs present in HSC+WT-EC-skin or WT-EC-skin+PBMC and KO-EC-skin+PBMC implanted skin grafts (**D)** 3D-printed skins with WT-EC but not KO-EC are rejected (inset of FACS hCD45 dot plot. **E)** Histopathologic evaluation of tissues show significant differences between HSC+skin and HSC+PBMC+KO-EC skin mice,in comparison with HSC+PBMC+WT-EC skin.

**Fig. 4.**
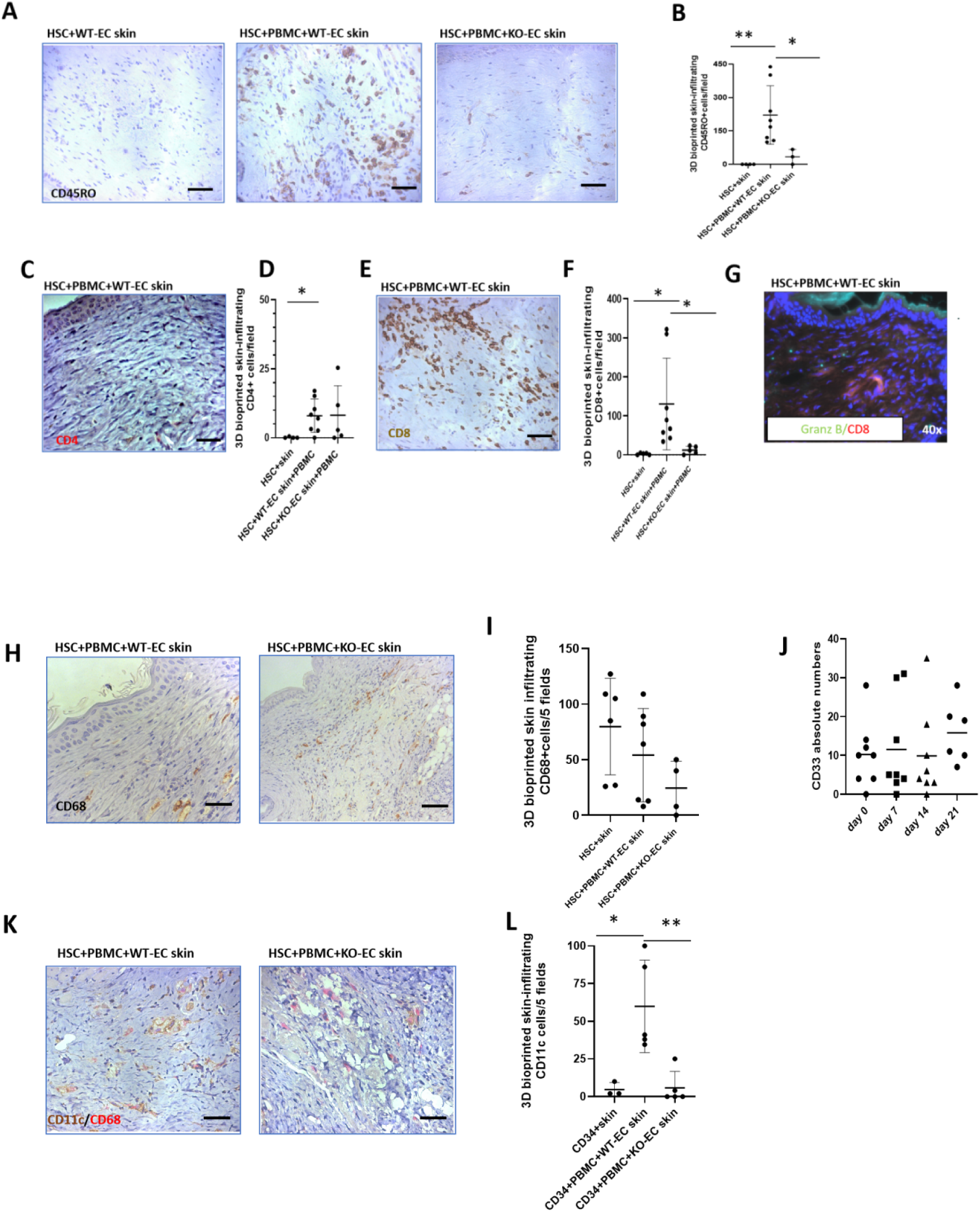
3D-printed skin-infiltrating cells. **A, B)** IHC staining of CD45RO shows massive expansion of memory T cells in WT-EC-skin+PBMC but not in KO-EC-skin+PBMC implanted skin grafts. **C, D)** Similar presence of CD4 lymphocytes found in WT+EC skin+PBMC or HSC+PBMC+KO-EC skin grafts. **E, F)** Massive expansion of CD8 lymphocytes described in HSC+PBMC+WT-EC skin group as compared to HSC+skin or HSC+PBMC+KO-EC skin cohort of animals. **G)** 3D-printed skin-infiltrating CD8 lymphocytes show Granzyme B positivity. **H, I)** IHC staining of CD68 shows infiltration of skin grafts by macrophages present in all 3 PBMC+skin, WT-EC-skin+PBMC and KO-EC-skin+PBMC implanted skin grafts. **J)** Stable circulating human myeloid cells described (CD33+ cells) at weekly assessment for 4 weeks. **K, L)** CD11c/CD68 double staining revealed infiltrating dendritic cells population in WT-EC but not KO-EC-skin mice.

We then focused on the nature of the human leukocytic infiltrates in the grafts containing WT-ECs. Consistent with our analysis of circulating cells, the majority of the infiltrate consisted of CD8+ T cells that expressed granzyme B (**Figure 4 C-G; CD4**: WT+EC skin+PBMC, n=8, 7.98±6.04 or HSC+PBMC+KO-EC skin, n=3, 8.16±10.72, HSC+skin, n=4, 0.125±0.25 vs HSC+PBMC+WT-EC skin, p=0.029; **CD8:** HSC+skin, n=4, 2.4±2.3 vs HSC+PBMC+WT-EC skin, n=8, 130.46±117.73, p=0.035). Such cells were absent in the grafts containing KO-ECs (**Figure 4 F, G;** HSC+PBMC+WT-EC skin vs HSC+PBMC+KO-EC skin, n=5, 12.18±10.06 p=0.049). Infiltrates in the rejecting grafts also included CD68+ macrophages and CD68-CD11c+ cells, which we tentatively identified as dendritic cells (**Figure 4 H, I, K, L, H, I; CD68:** PBMC+skin (n=6, 79.83±43.41), WT-EC-skin+PBMC (n=7, 54.14±41.69) and KO-EC-skin+PBMC (n=4, 24.5±24.24). Grafts with KO-EC cells showed infiltrating CD68+ macrophages but lacked CD11c+ cells (**Figure 4 H, I, K, L, CD11c:** HSC+skin, n=3, 4.66±4.62 vs HSC+PBMC+WT-EC skin, n=5, 59.9±30.7, p=0.028, HSC+PBMC+WT-EC skin vs HSC+PBMC+KO-EC skin, n=5, 5.8±10.87, p=0.0059) whereas mice receiving human HSCs, WT-ECs or KO-ECs skin grafts and human PBMCs showed stable numbers of circulating human myeloid cells followed weekly for 3 weeks after PBMC injection (**Fig 4 J**).

Finally, to assess the systemic nature of the immune response against the skin, we examined the spleens. The spleens of the animals with rejecting skin grafts were noticeably enlarged (0.8-1.1 cm vs 2.2-2.4 cm). Analysis of the spleens by both histology and flow cytometry confirmed the presence of hCD45+infiltration and large numbers of human CD8+ T cells in WT-EC skin grafts (**Figure 5 A, B, C, D; hCD45**: HSC+skin, n=6, 434±489.2 vs HSC+PBMC, n=3, 10.456.6±2738.98, p<0.0001, HSC+skin vs HSC+PBMC+WT-EC skinp=0.0005, HSC+PBMC+WT-EC skin, n=5, 10120.46±4456.34, vs HSC+PBMC+KO-EC skin, n=5, 1242.4±1485.17, p=0.0029, HSC+PBMC vs HSC+PBMC+KO-EC skin p=0.000, **CD8:** HSC+skin, n=5, 1±0.71 vs HSC+PBMC+WT-EC skin, n=6, 547.67±305.78, p=0.0033, HSC+PBMC+WT-EC skin vs HSC+PBMC+KO-EC skin, n=4, 115.25±133.84, p=0.03, HSC+skin vs HSC+PBMC, n=3, 391±180.02, p=0.0021).

**Fig. 5.**
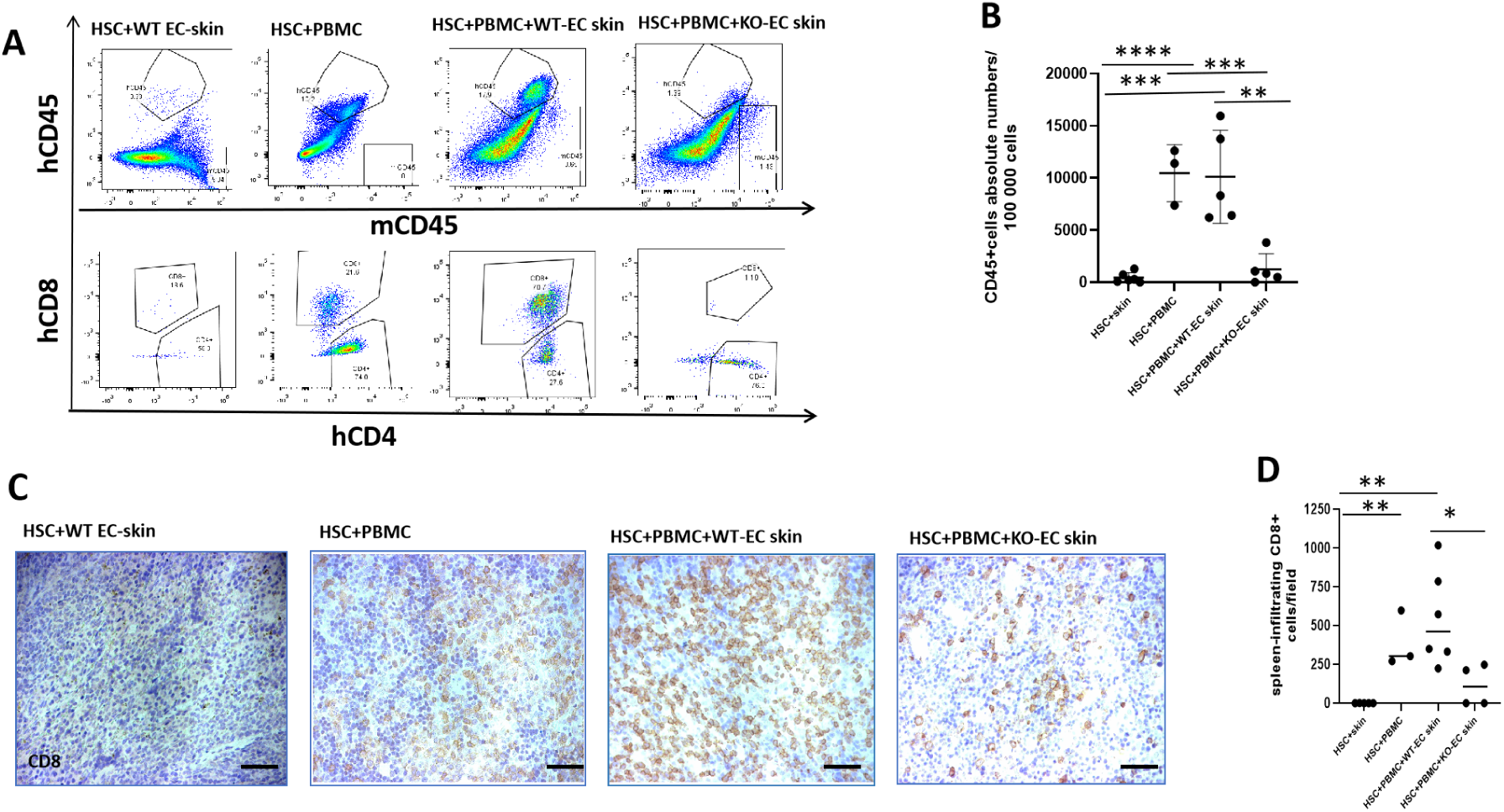
Characterization of spleen-infiltrating human cells. **A)** Comparison of infiltrating human cells by hCD45 and hCD4/CD8 staining. hCD45 staining show minimal hCD45 cells infiltration of spleens of HSC+PBMC+KO-EC skin animals. In consistency with circulation and skin grafts, robust expansion of CD8 lymphocytes is seen in HSC+PBMC+WT-EC skin animals whereas low infiltration is seen in HSC+PBMC+KO-EC skin mice. **B)** Comparison of circulating human leukocytes shows statistical significant hCD45 leukocytes of HSC+WT-EC skin+PBMC group as compared to HSC+skin or HSC+KO-EC skin+ PBMC groups. **C, D)** IHC staining reveals expansion of CD8 lymphocytes in HSC+PBMC+WT-EC skin mice.

Some animals receiving grafts with KO-ECs also showed splenic enlargement, but did not show expanded human CD8+ T cells.

Involvement of the spleen raised the issue of whether the CD8+ infiltrates were evidence of a systemic alloresponse against the graft or xenogeneic graft vs. host disease possibly triggered by the alloresponse. To address this question, we analyzed TCRβ sequences of the human T cells in the skin and spleens of 3 pairs of the mice showing evidence of graft rejection (all 3 from different donors cells). Strikingly, there was a large number of shared clones, including a few highly immunodominant clones in two of the three pairs, but the sequence of the immunodominant clones were not the same among the three animals (**Figure 6**). We interpret this as strong evidence for an alloresponse being systemic; the involvement of mouse tissues in addition to the skin graft is suggestive of semidirect recognition triggered by shedding of EC microparticles bearing intact MHC molecules into the circulation.

**Figure 6.**
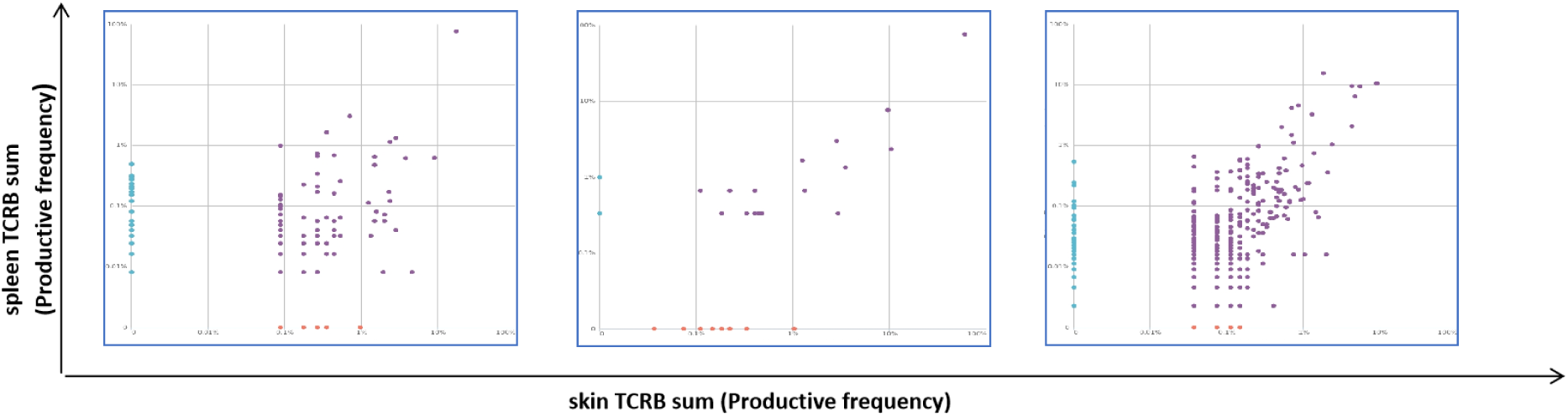
Immunosequencing of the T-Cell Receptor Repertoire of skin and matching spleen samples. High-throughput sequencing of the β chain of T cell receptors. X and Y axes show total amount of productive TCR β sequences in the skin or spleen samples with1-2 immunodominant clones in 2/3 pairs of investigated samples.

## DISCUSSION

Rodent models of transplantation are widely used for studying allograft rejection mechanisms, but many features of rodent immunology differ significantly from clinical experience. Human immune system mice appear to offer a possible bridge in which human cell-mediated rejection can be studied in a small animal model (*22*). Natural or synthetic human skin grafts in human immune system mice offer a way to analyze the role of blood vessels in the human allogeneic response. We and others have extensively exploited this approach, but an often overlooked limitation is the incompleteness of the engrafted human immune system.

Specifically, transplant rejection in the clinic not only involves direct T cell recognition of graft endothelium, but also involves innate immune cells, either as effector cells or as mediators of indirect or semi-direct recognition; these cells are missing in the most commonly used mouse models. Here, we have created a new model that addresses these prior limitations and reveals the importance of ECs in initiating rejection.

The MISTRG6 mouse has been engineered to allow development of innate myeloid and lymphocyte populations. As such, these mice are invaluable in studying responses to infection. Transplantation raises another issue, namely that a large element of the immune response to allografts is based on cross-reactive memory acquired from infections, and this cannot be easily generated in animals without significant fatality. We have addressed this problem by inoculated neonatal MISTRG6 mice with adult HSCs and then re-inoculating the same animals upon maturity with PBMCs, containing alloreactive effector memory T cells, from the same blood donor. Our exact protocol, developed from experience, is to place a skin graft three weeks before the administration of the PBMCs. This timing allows blood vessels in the graft to connect with host vessels, mimicking the establishment of surgical anastomoses in solid organ transplantation.

A second feature of our advanced model is that it uses synthetic rather than natural skin. This has two advantages. First, the knock-in of IL6 in the MISTRG6 model induces a non-physiological expansion of resident leukocytes in normal human skin that obscures transplant responses(*25*) . Second, and more importantly, it allows us to control the composition of the skin substitute. An important feature of our model is that the ECs, pericytes, fibroblasts and keratinocytes are all acquired from a single donor. To assemble these cell populations in a manner recapitulating key features of normal skin, we used 3D printing to separate the dermal from epidermal layer. This approach can be generalized for future work to incorporate other cell types, such as leukocytes, melanocytes or cells of skin adnexa.

There are two key findings of our study: First, that we can generate a rejection response that involves human myeloid cells as well as lymphocytes. Indeed, the myeloid compartment appears critical in triggering a T cell response, as judged by T cell expansion in the blood and spleen as well as the graft in animals receiving HSCs as well as PBMCs. The absence of myeloid cells at first seems to contradict our prior work in which adoptively transferred PBMC alone could cause rejection in C.B-17 SCID/bg mice. The simple explanation is that we had to introduce at least a hundred-fold more PBMCs or purified T cells to produce that result than we used in the current experiment. In the absence of myeloid cells, the T cells in the MISTRG6 animals are too few and simply fail to expand. Second, we show that it is EC presentation of antigen rather than increased vascular perfusion that leads to rejection. This conclusion has major implications for tissue engineering using allogeneic cells, namely that 1) ECs are a critical cell type for initiation of rejection and 2) MHC molecule expression on other cell types may be less critical. We note that this finding is consistent with the clinical experience using tissue engineered cellularized skin in which allogeneic fibroblasts and keratinocytes do not activate a recipient allogeneic rejection response.

While the current study offers a new model with significant advances compared to prior human immune system mouse studies, there are limitations. First, we do not know if ECs need to lack both class I or class II or both classes of MHC molecules to avoid rejection. Our prior work suggested class I molecules were more important, but both played a role (*20*).

Second, the systemic nature of the alloresponse in the present study raises the issue of shedding of intact MHC molecules and semi-direct presentation. This issue warrants further study. Third, while the protection from rejection when ECs lack both class I and class II MHC molecules appears fairly complete, we stopped the experiment at 3-4 weeks after PBMC inoculation because the mice that receive grafts with WT-ECs become fragile as circulating T cell numbers grow very large. In future work, we can assess if the animals receiving grafts containing ECs lacking MHC molecules—but where MHC expression is retained by other cell types-will reject their grafts eventually (by indirect presentation) or if modulation of EC immunogenicity is sufficient to eliminate rejection by tolerizing the immune response. Finally, we have not explored the role of alloantibody in graft rejection, an issue that also warrants further study. Despite these limitations, the present study significantly closes the gap between preclinical models and clinical experience. It also suggests that the recipient immune responses to tissue engineered grafts, like that to allogeneic organs is more dependent upon some cell types, such as ECs, than on others.

## MATERIALS AND METHODS

*Antibodies and other Reagents*: Antibodies used in these studies are described in **Supplementary Table 1**.

### Sex as a biological variable

Placenta, cord blood and foreskin served for cells isolations. Foreskin, as a source of fibroblasts and keratinocytes limited the option to include both female and male babies.

Both female and male mice were utilized in the 3D-printed-skin implantation experiments with no differences observed.

### 3D-printed skin graft preparation

Anonymized tissue donors for 3D bioprinted skin production were acquired through the Yale University Reproductive Sciences Biobank from discarded tissues with informed consent. Cells involved in the skin graft printing were isolated, cultured and characterized as described, as were the composition of the bioinks the 3D printing process and surgical engraftment (*14*). In brief, human pericytes were isolated from discarded placenta, endothelial colony forming cells (ECFCs) were isolated from cord blood, fibroblasts and keratinocytes were isolated from discarded male foreskins and each cell type was expanded in vitro. All four cell types used in each graft were obtained from a single male infant donor and a total of 3 different donors were used over the course of these studies and gave similar results. Where indicated, unmodified ECFC-derived ECs (WT-ECs) were replaced with ECFC-derived ECs from the same donor that had been modified by CRISPR/Cas9 as described to ablate expression of class I and II MHC molecules (*20*). 3D bioprinting was performed with a BioX bioprinter (CELLINK) as described. The only change in the process from our original description was that the dermal bioink was introduced into a sterile nonwoven polyglycolic acid mesh (Confluent Medical Technologies, Inc) to delay contraction after implantation, allowing the dermal cells more time to synthesize extracellular matrix. As we described previously, the mesh is absorbed by three weeks in the absence of human leukocytes with minimal inflammation and no effect on graft cell viability (*26*).

### Animals

MISTRG6 mice were engineered by a human/mouse homolog gene-replacement strategy to provide physiological expression with regard to quantity, location and time of M-CSF (monocytes and tissue macrophage development), GM-CSF/IL-3 (lung alveolar macrophages), SIRPα (tolerance of macrophages to human cells), ThPO (hematopoiesis and platelets) and IL-6 (improved engraftment and antibody responses), in a Rag2/Gamma common chain deleted background, were generated (*22, 23*). All mice were maintained under specific pathogen free conditions in Yale School of Medicine animal facilities. Availability of mice was limiting and both males and females were used interchangeably with no obvious differences in outcomes.

### Experimental outline

The general experimental protocol used was described in **Figure 1**. MISTRG 6 neonates were irradiated, anti-CD3 antibody-treated and injected intrahepatically within 24-48 hrs after birth with human adult CD34+ hematopoietic stem cells (HSCs) stem cells obtained from a cryopreserved, discarded and anonymized GM-CSF-mobilized donor leukapheresis collection. HSCs were isolated by density gradient centrifugation (Lymphoprep, StemCell Technologies) followed by immunomagnetic selection (EasySep Human CD34 Positive Selection Kit, StemCell Technologies) and assessed by flow cytometry for purity. Isolations contained over 93% CD34+ cells. Once the animals showed stable levels of circulating human cells, confirmed by flow cytometry (typically at 10 weeks) 3D bioprinted skin grafts, containing cells allogeneic to injected CD34+ cells, were implanted on dorsal part of MISTRG6 mice. Healing in of the skin grafts after surgery was allowed to occur for 2 weeks. At that point, mice were injected with autologous PBMCs to the HSCs. Mice were monitored for signs of rejection and harvested at various time points (Day 17, Day 23, Day 28-34). Control groups in some experiments omitted HSC injection or omitted PBMC injection or omitted skin grafting, or were left uninjected, described as follows (1) mice receiving human adult HSCs as neonates and human PBMCs from the same individual as adults without skin grafting; (2) mice receiving a 3D-printed human skin graft made with WT-ECs as adults in the absence of human hematopoietic cells; (3) mice receiving human HSCs as neonates and a 3D-printed human skin with WT-ECs from a source allogeneic to the HSCs as adults without a PBMC boost; (4) mice receiving human HSCs as neonates and a 3D-printed human skin with KO-ECs from a source allogeneic to the HSCs as adults; and (5) mice receiving 3D-printed human skin with WT-ECs as adults and PBMCs from a donor allogeneic to the skin cell source 3 weeks later but without neonatal inoculation with human HSCs. Groups 1-5 constitute our controls and key findings are summarized in **Supplementary Table 2 and Supplementary Figure 2**. Overall, we analyzed 47 MISTRG6 mice using one leukocyte donor and 3 skin donor collections. Not every experiment contained all of the controls as we were limited by litter size and availability of MISTRG6 animals.

### Flow cytometry

Flow cytometry was performed on LSR2 or Fortessa instruments (both Becton Dickinson) to characterize cell populations used for 3D bioprinting (as described in ref 14), and for characterization of circulating cells and cells extracted from tissue samples at the time of sacrifice. Circulating cells were assessed for species origin (using mouse and human CD45) as well as for human cell types (CD3 T cells, CD19 B cells, NKp46 NK cells, CD33 (myeloid cells) throughout the experiment as indicated in Figure 1. Cells extracted from skin grafts and from spleens were analyzed at the time of sacrifice for CD3 T cells and subpopulations (CD4 and CD8) effector memory (loss of CCR7). CD8+ T cells were additionally assessed for Granzyme B indicative of cytolytic potential. Flow cytometric data were analyzed by Flow Jo software (BD Biosciences).

### Morphological Analysis of Tissue Samples

Hematoxylin and eosin (H&E) staining, immunohistochemical and immunofluorescent staining were performed on flash-frozen cryosections and formalin-fixed-paraffin embedded (FFPE) sections as indicated in each panel (*14*). The H&E stained sections were used to determine the extent of rejection in skin samples (day 17, day 23, 28-34) were assessed. Histopathological score of skin transplantation rejection was determined based on the following scoring system: 0-as no graft inflammation, normal vessels and endothelial cells lining, 1-as infiltrating cells (inflammation), no vascular damage, 2- as infiltrating cells (inflammation) and signs of vascular damage and 3- as infiltrating cells (inflammation) and profound vascular damage, assessed by a dermatopathologist JMM) blinded to the treatment groups.

The presence and viability of bioprinted and implanted dermal graft cells were assessed by HLA-B staining. Skin grafts were evaluated for markers as CD56, CD68, CD11c, CD45RO, human and mouse CD31, CD3, CD4, CD8, Granzyme B, or E-Selectin. All immunofluorescent samples were analyzed by EVOS (Thermo) or confocal microscopy (Stellaris 5, Leica) and IHC samples by Zeiss Axiovert (Supplementary Table 1, List of antibodies).

### TCR sequencing

Comprehensive analysis of TCR receptor b chain was performed by Adaptive Biotechnologies (Seattle, WA) to assess the clonality of infiltrating lymphocytes in skin grafts and spleens of 3 different skin donors, all with the same HSC and PBMC donor. (FFPE curls, 50 um). The sequencing repertoire data were analysed by immunoSEQ® 3.0 Analyzer (Adaptive Biotechnologies) and a freely available VDJTools software package for programming language R (*27*).

### Statistical analysis

Statistical analysis was performed by using 2-tailed unpaired t test by GraphPad Prism 10.3.0 software. The data are presented as mean±SD. Differences with a value of P<0.05 was considered as statistically significant. Means, standard deviations and p values are described in the Results section for each group.

### Study approval

Deidentified human tissues for 3D bioprinted skin production were acquired through the Yale University Reproductive Sciences Biobank from discarded tissues with informed consent.

Mice experiments were performed in compliance with approved Institutional Animal Care and Use Committee (IACUC) protocol.

### Data availability statement

All data supporting the findings of this study will be available online at https://datadryad.org

## Acknowledgement

This work was supported by 1R01HL169238-01A1.

We thank Dr. Diane Krause for providing cells used in the study, Amos Brooks and Yale Pathology Tissue Services for their assistance with FFPE tissue samples. We also thank Ewa Menet, Yale Flow Cytometry Core for her assistance with modified endothelial cells sorting. The Yale Flow Cytometry Core is supported in part by an NCI Cancer Center Support Grant P30 CA016359. We thank families consenting for cord blood, placenta and foreskin matching tissue donations. Tissue acquisitions were supported by the Yale University Reproductive Sciences Biobank (HIC#12696), a component of the Department of Obstetrics, Gynecology & Reproductive Sciences, Yale School of Medicine, New Haven, CT.

## Authors contributions

Concept and Design: ZT, WMS, JSP. Acquisition of materials, annotation and interpretation of data: all authors. Data analysis: ZT, JMM, JSP. Drafting of the manuscript: ZT, WMS, JSP. Critical editing of the manuscript: all authors.

---

**Supplementary Figure 1.**
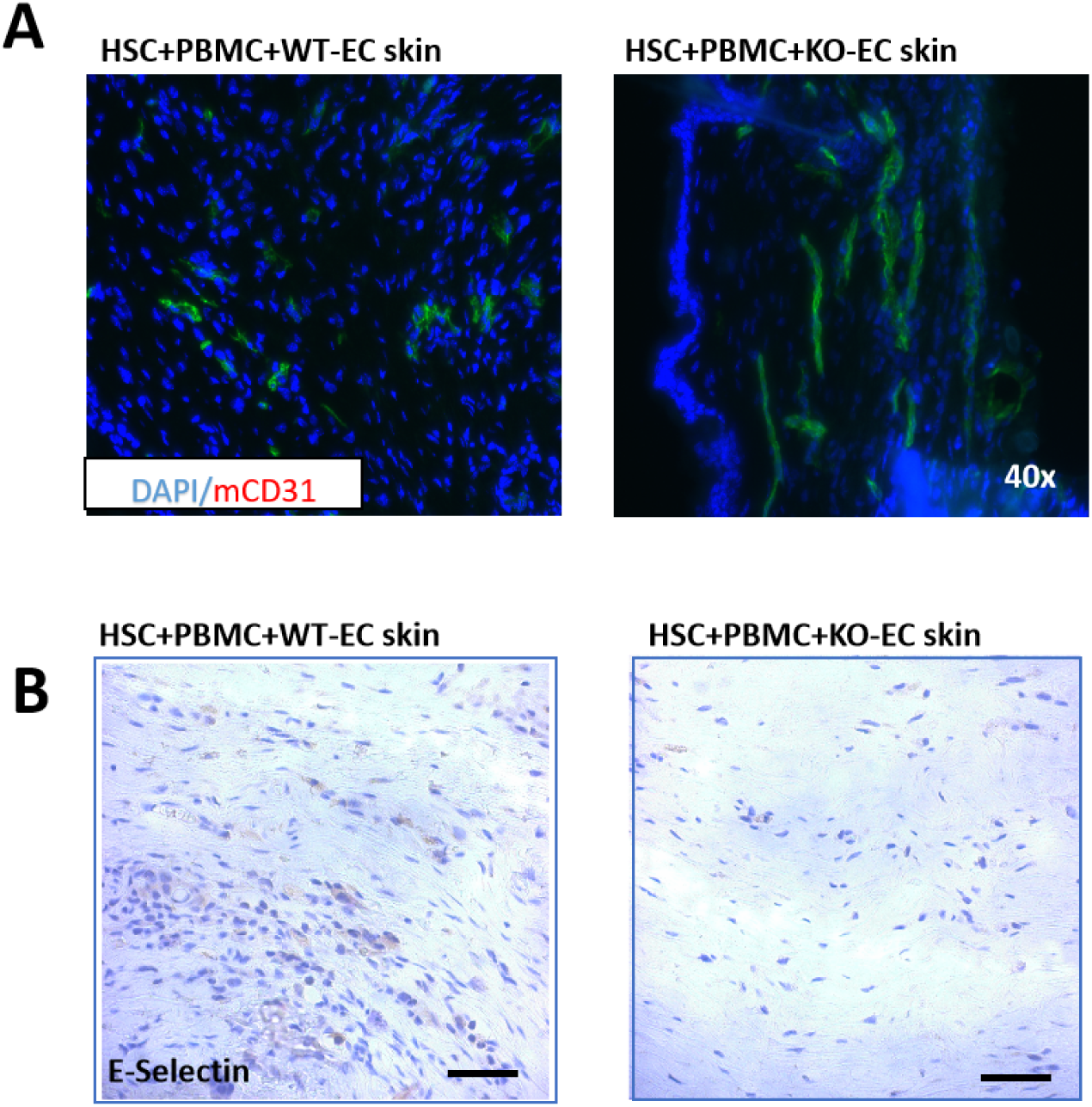
Further characterization of implanted 3D-printed skin grafts. **A)** Staining of mCD31 infiltrating human part of skin shows similar presence of mice endothelial cells in the HSC+PBMC+WT-EC skin or HSC+PBMC+KO-EC skin grafts indicating undamaged mouse endothelial cells. **B)** Presence of E-Selectin staining in HSC+PBMC+WT-EC tissues whereas no activation marker positivity in HSC+PBMC+KO-EC skin grafts.

**Supplementary Figure 2.**
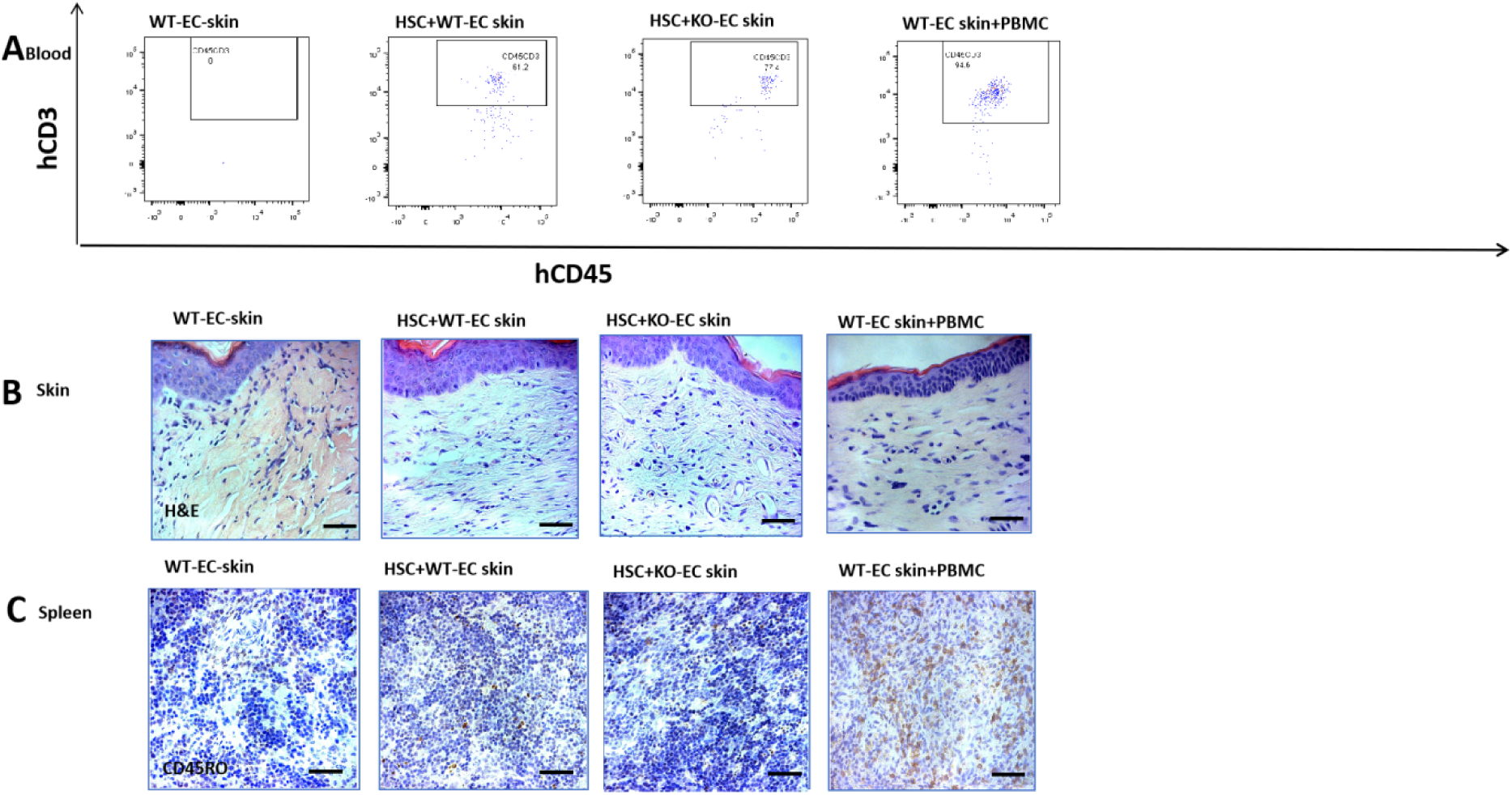
Blood, skin and spleen samples of control groups (2)-(5) (described in details in the Methods section). **A)** Dot plots of circulating human CD45/CD3 cells in (2) WT-EC skin mice, (3) HSC+WT-EC skin mice, (4) HSC+KO-EC skin mice and (5) WT-EC-skin+PBMC mice. **B)** H&E staining of 3D-printed skins grafts of (2) WT-EC skin mice, (3) HSC+WT-EC skin mice, (4) HSC+KO-EC skin mice and (5) WT-EC-skin+PBMC mice. **C)** CD45RO staining of spleens of (2) WT-EC skin mice, (3) HSC+WT-EC skin mice, (4) HSC+KO-EC skin mice, (5) WT-EC-skin+PBMC mice indicating infiltration of spleens by human memory T cells in WT-EC-skin+PBMC mice only.

**Supplementary Table 1.**
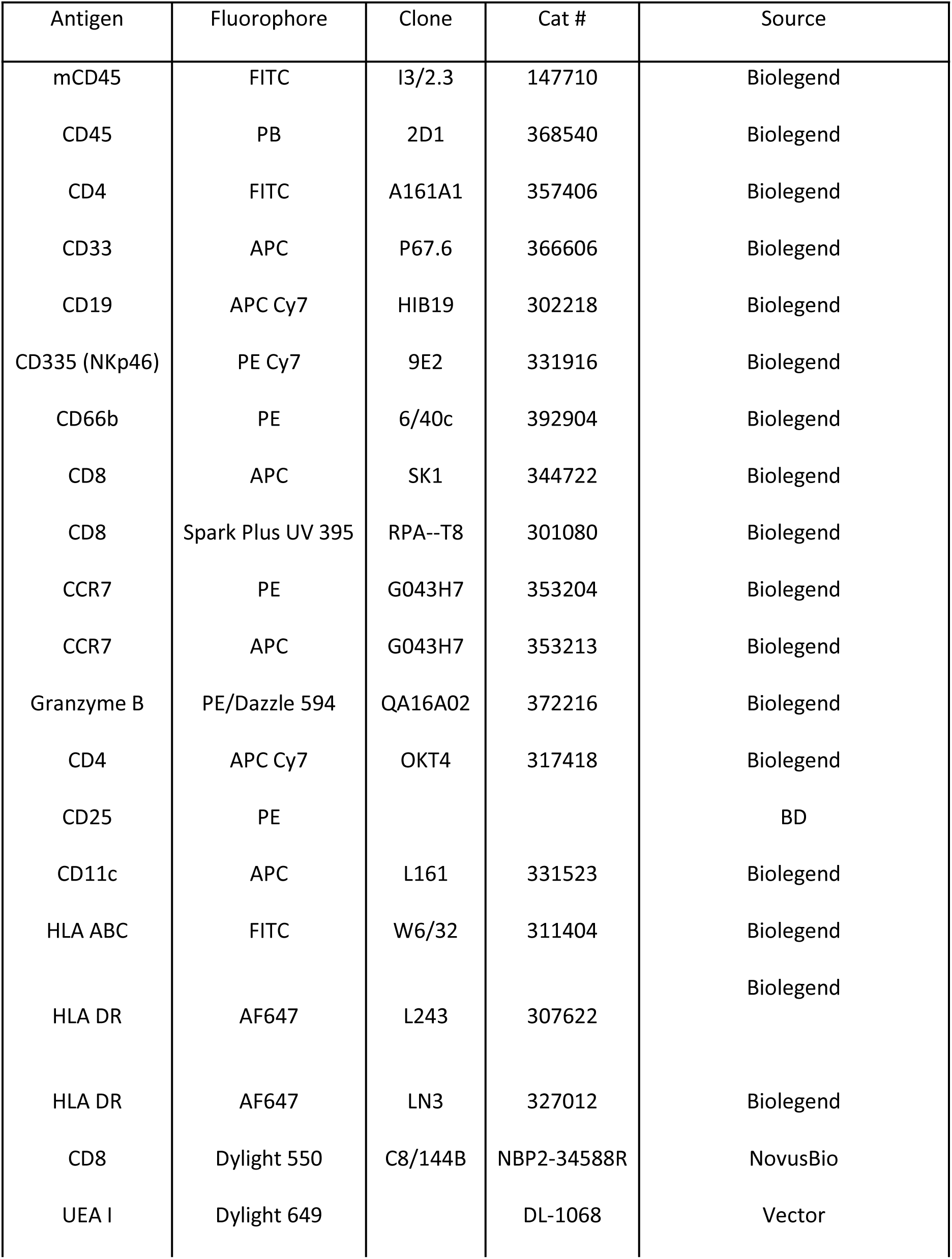
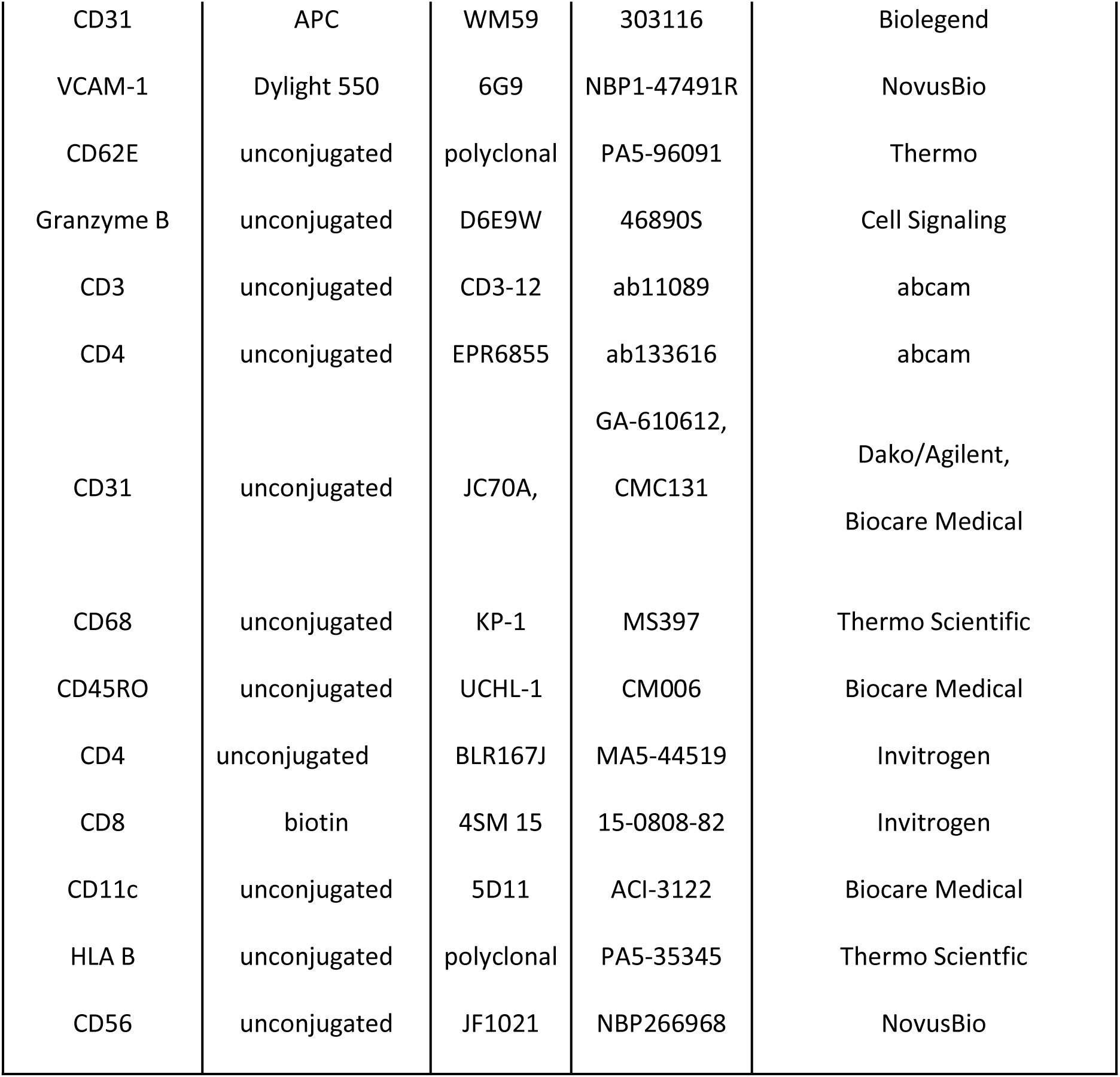
List of used antibodies.

**Supplementary Table 2.**
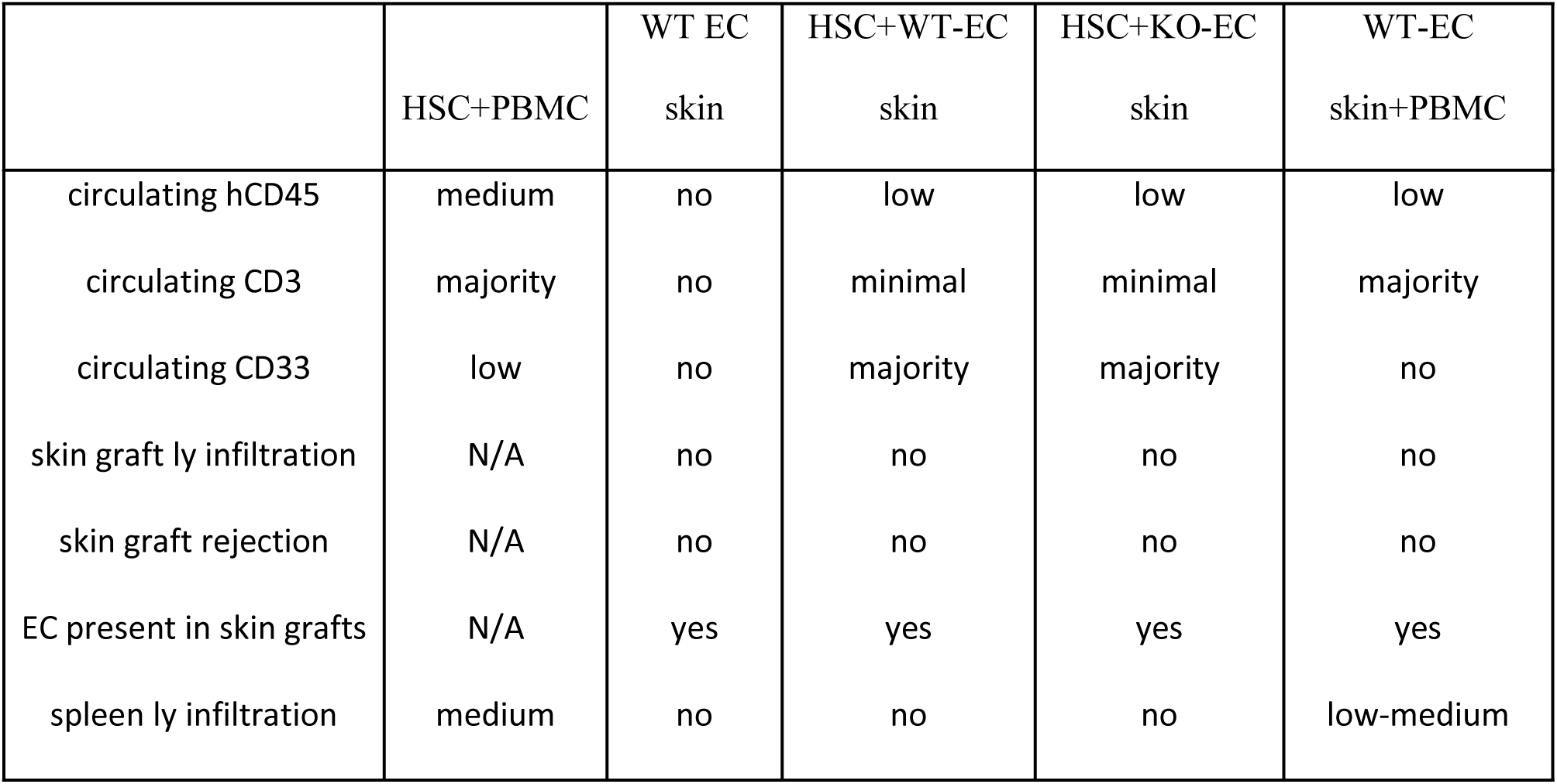
Assessed parameters of control groups.

