## Supplementary Figures and Tables for "Endothelial cell MHC molecules are necessary and sufficient to reject 3D-printed human skin grafts in an advanced human immune system mouse"

**Supplementary Figure 1.**

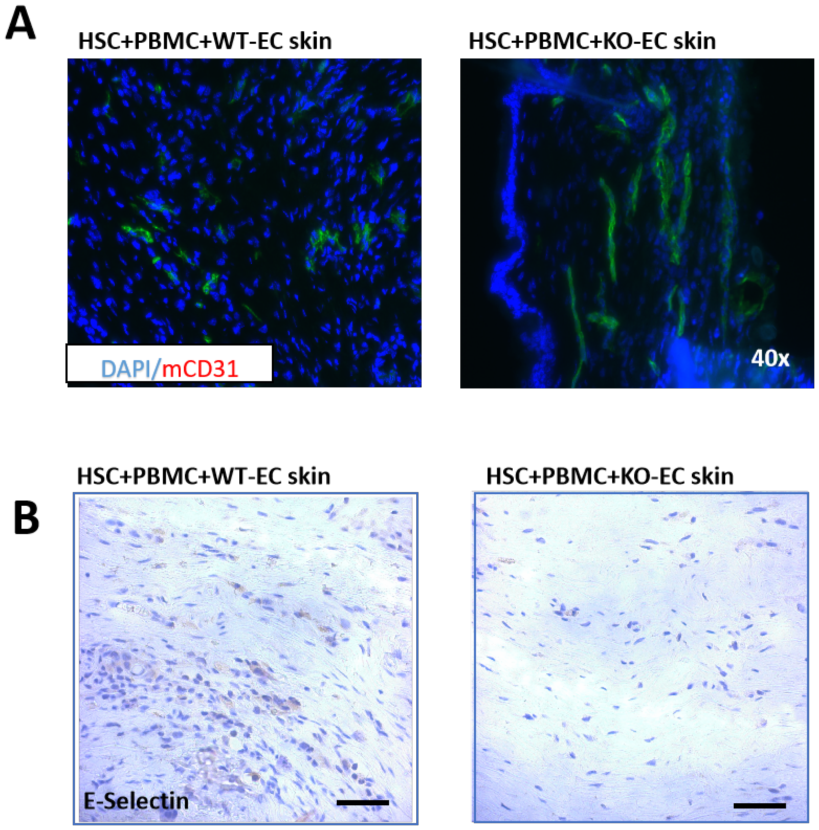


**Supplementary Figure 1. Further characterization of implanted 3D-printed skin grafts**

**A)** Staining of mCD31 infiltrating human part of skin shows similar presence of mice endothelial cells in the HSC+PBMC+WT-EC skin or HSC+PBMC+KO-EC skin grafts indicating undamaged mouse endothelial cells.

**B)** Presence of E-Selectin staining in HSC+PBMC+WT-EC tissues whereas no activation marker positivity in HSC+PBMC+KO-EC skin grafts.

**Supplementary Figure 2.**

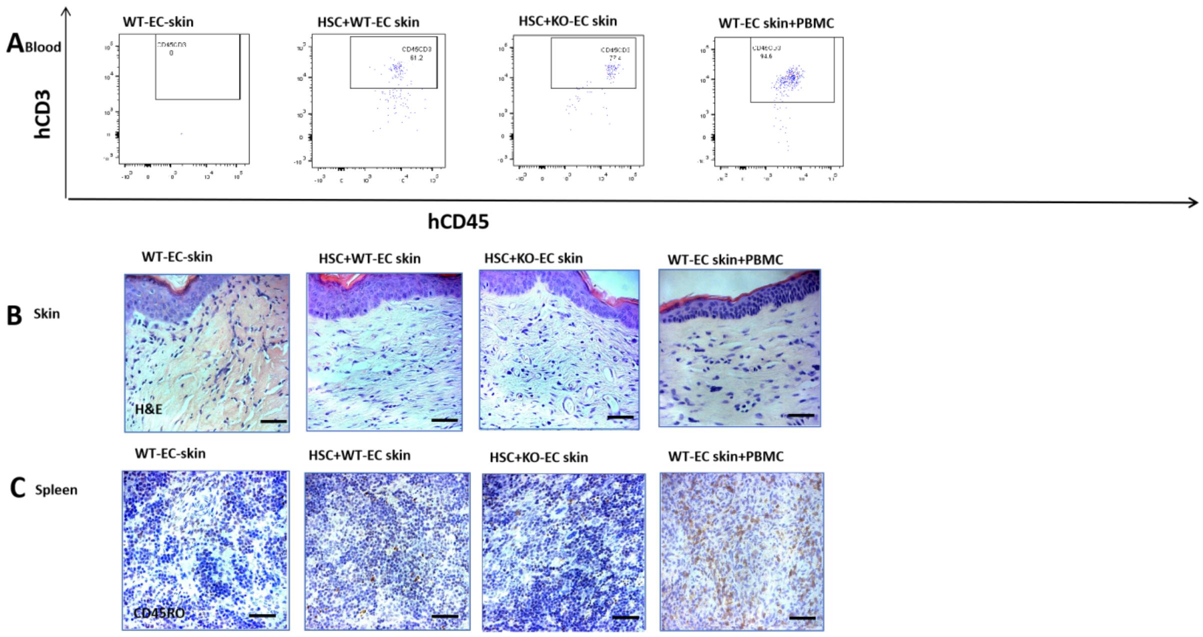


**Supplementary Figure 2. Blood, skin and spleen samples of control groups (2)-(5) (described in details in the Methods section).**

**A)** Dot plots of circulating human CD45/CD3 cells in (2) WT-EC skin mice, (3) HSC+WT-EC skin mice, (4) HSC+KO-EC skin mice and (5) WT-EC-skin+PBMC mice.

**A)** H&E staining of 3D-printed skins grafts of (2) WT-EC skin mice, (3) HSC+WT-EC skin mice, (4) HSC+KO-EC skin mice and (5) WT-EC-skin+PBMC mice.

**B)** CD45RO staining of spleens of (2) WT-EC skin mice, (3) HSC+WT-EC skin mice, (4) HSC+KO-EC skin mice, (5) WT-EC-skin+PBMC mice indicating infiltration of spleens by human memory T cells in WT-EC-skin+PBMC mice only.

**Supplementary Table 1. List of used antibodies**

| Antigen | Fluorophore | Clone | Cat # | Source |
|---|---|---|---|---|
| mCD45 | FITC | I3/2.3 | 147710 | Biolegend |
| CD45 | PB | 2D1 | 368540 | Biolegend |
| CD4 | FITC | A161A1 | 357406 | Biolegend |
| CD33 | APC | P67.6 | 366606 | Biolegend |
| CD19 | APC Cy7 | HIB19 | 302218 | Biolegend |
| CD335 (NKp46) | PE Cy7 | 9E2 | 331916 | Biolegend |
| CD66b | PE | 6/40c | 392904 | Biolegend |
| CD8 | APC | SK1 | 344722 | Biolegend |
| CD8 | Spark Plus UV 395 | RPA--T8 | 301080 | Biolegend |
| CCR7 | PE | G043H7 | 353204 | Biolegend |
| CCR7 | APC | G043H7 | 353213 | Biolegend |
| Granzyme B | PE/Dazzle 594 | QA16A02 | 372216 | Biolegend |
| CD4 | APC Cy7 | OKT4 | 317418 | Biolegend |
| CD25 | PE | | | BD |
| CD11c | APC | L161 | 331523 | Biolegend |
| HLA ABC | FITC | W6/32 | 311404 | Biolegend |
| HLA DR | AF647 | L243 | 307622 | Biolegend |
| HLA DR | AF647 | LN3 | 327012 | Biolegend |
| CD8 | Dylight 550 | C8/144B | NBP2-34588R | NovusBio |
| UEA I | Dylight 649 | | DL-1068 | Vector |
| CD31 | APC | WM59 | 303116 | Biolegend |
| VCAM-1 | Dylight 550 | 6G9 | NBP1-47491R | NovusBio |

| | | | | |
|---|---|---|---|---|
| CD62E | unconjugated | polyclonal | PA5-96091 | Thermo |
| Granzyme B | unconjugated | D6E9W | 46890S | Cell Signaling |
| CD3 | unconjugated | CD3-12 | ab11089 | abcam |
| CD4 | unconjugated | EPR6855 | ab133616 | abcam |
| CD31 | unconjugated | JC70A, | GA-610612, CMC131 | Dako/Agilent, Biocare Medical |
| CD68 | unconjugated | KP-1 | MS397 | Thermo Scientific |
| CD45RO | unconjugated | UCHL-1 | CM006 | Biocare Medical |
| CD4 | unconjugated | BLR167J | MA5-44519 | Invitrogen |
| CD8 | biotin | 4SM 15 | 15-0808-82 | Invitrogen |
| CD11c | unconjugated | 5D11 | ACI-3122 | Biocare Medical |
| HLA B | unconjugated | polyclonal | PA5-35345 | Thermo Scientfic |
| CD56 | unconjugated | JF1021 | NBP266968 | NovusBio |

**Supplementary Table 2. Assessed parameters of control groups**

| | HSC+PBMC | WT EC skin | HSC+WT-EC skin | HSC+KO-EC skin | WT-EC skin+PBMC |
|---|---|---|---|---|---|
| circulating hCD45 | medium | no | low | low | low |
| circulating CD3 | majority | no | minimal | minimal | majority |
| circulating CD33 | low | no | majority | majority | no |
| skin graft ly infiltration | N/A | no | no | no | no |

| | | | | | |
|---|---|---|---|---|---|
| skin graft rejection | N/A | no | no | no | no |
| EC present in skin grafts | N/A | yes | yes | yes | yes |
| spleen ly infiltration | medium | no | no | no | low-medium |
